# Comparative phylogeography of Ponto-Caspian amphipods throughout the invaded Baltic and native NW Black Sea donor ranges – can introduction mode affect genetic diversity?

**DOI:** 10.1101/2023.01.18.524525

**Authors:** Denis Copilaș-Ciocianu, Eglė Šidagytė-Copilas, Mikhail O. Son, Halyna Morhun, Jan Niklas Macher, Kęstutis Arbačiauskas

## Abstract

The Baltic countries harbor a diverse assemblage of alien amphipods of Ponto-Caspian origin. The composition of this fauna was shaped by three invasion waves: 1) pre-20^th^ century dispersals via watershed-connecting canals, 2) deliberate introductions in the 1960s, and 3) new dispersals during the last decade via shipping and pre-existing canals. Given this rich invasion history, we test whether genetic diversity (mitochondrial and nuclear) differs between the native and invaded ranges and between the deliberately introduced species and the ones that dispersed on their own. Our results show a significant decrease in mitochondrial but not nuclear genetic diversity in the invaded Baltic range. We also find that in the invaded range the introduced species exhibit a higher mitochondrial and nuclear genetic diversity than the species that dispersed on their own, while in the native range only the nuclear diversity is higher in introduced species. Mitochondrial diversity was more structured geographically in the native range and the dominant invasive haplotypes were detected in the native populations of all but one species, further highlighting the utility of this marker in tracing invasion sources. Our comparative approach provides important insight into the inter-range genetic diversity of Ponto-Caspian invaders, highlighting the role of introduction mode.

## Introduction

The importance of genetic variation on the outcome of biological invasions has been recognized for decades (Baker & Stebbins, 1965). Initially, it was thought that small founding alien populations would be subjected to strong genetic drift and inbreeding, leading to severe loss of genetic diversity and hampering adaptation to the novel environment by reducing fitness and evolutionary potential (Estoup et al., 2016). However, it later became apparent that the loss of genetic diversity is not so prevalent since the effects of bottlenecks can be overruled by various factors such as high propagule pressure, admixture between invasive populations of different origin, and spatially structured genetic diversity of source populations (Kolbe et al., 2007; Roman & Darling, 2007). Importantly, loss of variation at the commonly employed selectively neutral genetic markers does not necessarily entail a reduction in variation at ecologically relevant traits that are under selection (Dlugosch et al., 2015). Therefore, the so-called “genetic paradox of biological invasion” (i.e. successful adaptation without genetic variation) is valid only for a few species (Estoup et al., 2016).

The Ponto-Caspian region has long been recognized as one of the most significant source of aquatic invasive species throughout the Northern Hemisphere (Bij de Vaate et al., 2002; Copilaș-Ciocianu et al., 2022b). The fauna of this area is ecologically diverse, adaptable and tolerant to large salinity fluctuations, making it particularly successful at colonizing and rapidly multiplying in new habitats (Reid & Orlova, 2002; Arbačiauskas et al., 2013; Šidagytė & Arbačiauskas, 2016; Hupało et al., 2018; Meßner & Zettler, 2018; Paiva et al., 2018; Cuthbert et al., 2020; Copilaș-Ciocianu & Sidorov, 2022). As such, Ponto-Caspian invasions are generally associated with significant ecological and sometimes economic damage (Vanderploeg et al., 2002; Strayer, 2009; Haubrock et al., 2022). Among this melting pot of Ponto-Caspian invaders, amphipod crustaceans seem to be the most numerous, with up to 40% of the entire fauna expanding beyond the native range (Copilaș-Ciocianu et al., 2022b), often with fatal consequences for the native species (Dermott et al., 1998; Arbačiauskas, 2008; Grabowski et al., 2009; Rewicz et al., 2014; Soto et al., 2022).

The Baltic region and specifically Lithuania have a particularly rich history of Ponto-Caspian amphipod invasions (Arbačiauskas et al., 2011). The first invasion wave took place well before the 20^th^ century and was enabled by the construction of artificial canals that connected the Baltic and Black Sea watersheds, providing a dispersal corridor for Ponto-Caspian species. Through this first wave *Chelicorophium curvispinum* (Sars, 1895) and *Chaetogammarus ischnus* (Stebbing, 1899) reached the area (Jarocki & Demianowicz, 1931; Jażdżewski, 1980), although the latter does not occur there anymore (Arbačiauskas et al., 2017; Copilaș-Ciocianu et al., 2022b). The second and most important wave consisted of intentional introductions in the 1960s with the aim of improving fish feed (Gasiūnas, 1963). Several peracarid species, including four amphipods (*C. ischnus, C. warpachowskyi* (Sars, 1894), *Obesogammarus crassus* (Sars, 1894), and *Pontogammarus robustoides* (Sars, 1894)) were initially introduced and acclimated in the Kaunas Water Reservoir (WR) in Lithuania (though *C. ischnus* later went extinct) (Jażdżewski, 1980; Vaitonis et al., 1990; Arbačiauskas et al., 2017). From there they either were deliberately spread to other waterbodies and neighboring countries (until the late 1980s) or dispersed on their own throughout most of the Baltic basin and beyond (Arbačiauskas et al., 2011; Moedt & van Haaren, 2018; Meßner & Zettler, 2021). The sources of these species’ translocations were the then newly-built Dnieper and Simferopol WRs in Ukraine which were artificially populated with specimens originating from the native Dnieper-Bug estuary (Arbačiauskas et al., 2017). The third and last invasion wave brought two more species (*Dikerogammarus haemobaphes* (Eichwald, 1841) and *D. villosus* (Sowinsky, 1894)) in the last decade which dispersed on their own via the previously built canals or the Baltic Sea (Šidagytė et al., 2017; Copilaș-Ciocianu & Šidagytė-Copilas, 2022). On-going regional expansion is continuously reported in all of these species throughout the Baltic region (Grudule et al., 2007; Arbačiauskas et al., 2017; Minchin et al., 2019; Lipinskaya et al., 2021; Copilaș-Ciocianu & Šidagytė-Copilas, 2022).

The diverse history of introductions of invasive Ponto-Caspian species to the Baltic region makes this area an interesting model system for comparative studies on the genetic diversity of closely related invaders and how it is influenced by invasion history. As such, with this paper we aim to answer two main questions outlined below.

**Question 1**: Do invasive populations exhibit a decrease in genetic diversity relative to the donor populations? Considering the adaptability and success of Ponto-Caspian species in non-native areas, one could assume that genetic diversity is not substantially reduced. Indeed, recent studies have shown that invasive populations of multiple Ponto-Caspian species show comparable genetic diversity with the native populations, especially at the nuclear genome (Stepien et al., 2005; Rewicz et al., 2015; Audzijonyte et al., 2017; Jażdżewska et al., 2020), although this is not always the case (Cristescu et al., 2001, 2004; Rewicz et al., 2017).

**Question 2**: Is there a difference in genetic diversity patterns between the deliberately introduced species (i.e. *C. warpachowskyi, O. crassus, P. robustoides*) and the ones that dispersed on their own (i.e. *C. curvispinum, D. villosus, D. haemobaphes*)? Since multiple factors influence the genetic diversity of invasive populations, we may expect noticeable differences between the introduced and dispersed species. On one hand, the deliberately introduced species were released in high numbers (thousands) of individuals at once, possibly retaining a significant proportion of the original genetic diversity due to a less stringent effect of genetic drift. Contrastingly, species that spread on their own are on the northern limit of their invaded range in the Baltic area (Copilaș-Ciocianu et al., 2022b) and are possibly under stronger selective pressure due to prolonged dispersal along an increasing gradient of environmental harshness. Given that these factors are known to reduce genetic diversity (Hardie & Hutchings, 2010; Colautti & Lau, 2015), one could expect that the species that arrived via dispersal would have a reduced genetic diversity in comparison to the deliberately introduced species. On the other hand, species arriving via dispersal could harbor significant genetic diversity due to a higher propagule pressure than the introduced species which were transplanted only once from the native region (Roman & Darling, 2007).

As such, examining the genetic diversity of the invasive Ponto-Caspian amphipods among ranges and introduction modes could provide important insight into their long-term persistence, highlight their adaptation and evolutionary potential, and confirm their geographical origin.

## Materials and methods

### Sampling

Six species were targeted: three deliberately introduced in the Baltic region (*C. warpachowskyi, O. crassus*, and *P. robustoides*) and three that dispersed on their own to this region (*C. curvispinum, D. haemobaphes*, and *D. villosus*). The sampling was designed to thoroughly cover both the native donor range (lagoons and estuaries throughout the NW Black Sea coast—26 sites) and the invaded Baltic range (lagoons, rivers and lakes belonging to the SE Baltic Sea drainages—37 sites). In the native region we sampled specifically the lagoons and estuaries (Bug-Dnieper) which were the initial sources, as well as the Simferopol WR in Ukraine which was artificially populated with Dnieper-Bug specimens and from where the deliberately introduced species we transplanted to Lithuania. Unfortunately, despite intense effort no amphipods were sampled from the Dnieper WR, which was also artificially populated with Bug-Dnieper specimens that were subsequently introduced to Lithuania. Adjacent regions in Romania and Bulgaria were also sampled in order to gain a better understanding of the regional genetic diversity and to pinpoint the source populations of the three species that dispersed on their own (Table S1, Fig. 1). In the invaded Baltic region the sampling covered Estonia, Latvia and Lithuania with a special focus on the latter since it was the epicenter of introductions. Additionally, we also included two sites from Poland (Vistula and Szczecin lagoons) as these were also invaded from the Baltic countries by two of the deliberately introduced species (*O. crassus*, and *P. robustoides*).

**Fig. 1.**
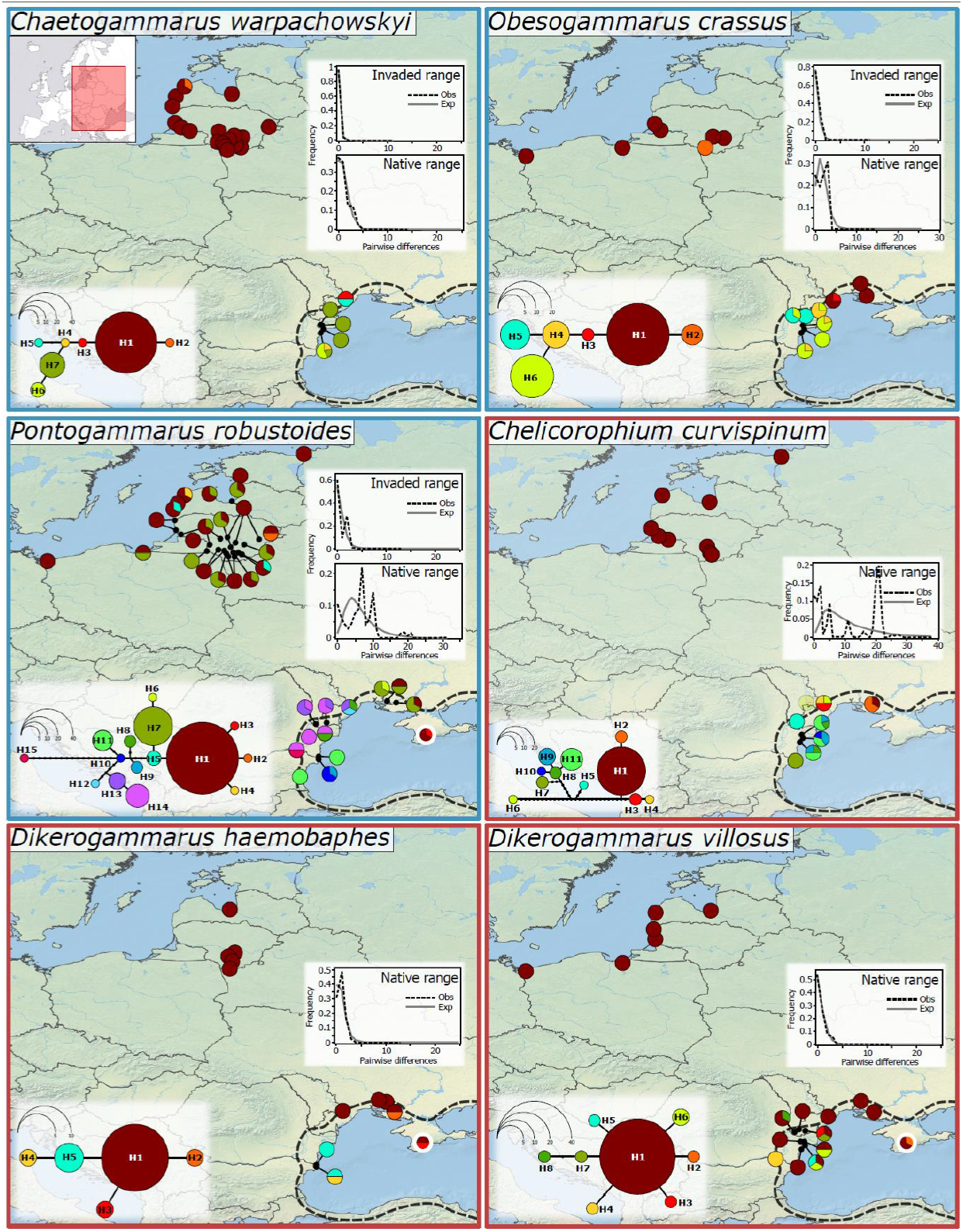
Patterns of mitochondrial (COI) haplotype distribution between the native and invaded ranges. Insets on the lower left depict haplotype networks while on the upper right are mismatch distributions. The native range is shown with a dashed black line. The site indicated with a thick white outline is the Simferopol WR which lies in the native range but was artificially populated. Species that were deliberately introduced and that dispersed on their own are indicated with a blue and red frame, respectively.

Animals were sampled in the late summer/early autumn in 2012, 2020, and 2021 (Table S1). All possible habitats were sampled along shorelines in shallow water up to 1.5 m depth using kick sampling with a hand net. Specimens were preserved in the field in 96% ethanol which was replaced several times. In the laboratory the material was stored at −20°C in fresh ethanol. Specimens were identified under a stereomicroscope using the latest keys (Copilaș-Ciocianu & Sidorov, 2022)

### Laboratory protocols

Genomic DNA was extracted as described in Copilaș-Ciocianu et al. (2022). Briefly, a piece of dorsal tissue was excised using a biopsy punch and DNA was isolated using the Quick-DNA Miniprep Plus Kit (Zymo Research). Depending on the available material, between one and five individuals per sampling location were used for genetic analyses. Two protein-coding markers were chosen for sequencing: the mitochondrial cytochrome *c* oxidase subunit one (COI) and the nuclear long-wave opsin (OPS). Previous studies indicated that these makers have sufficient variation to track invasion pathways and explore genetic diversity of invasive Ponto-Caspian crustaceans (Audzijonyte et al., 2008, 2017; Rewicz et al., 2015; Morhun et al., 2022). For *C. warpachowskyi* sequencing of OPS failed and the nuclear large ribosomal subunit (28S) was sequenced instead which has a comparable level of variation. The PCR protocols for COI followed Copilaș-Ciocianu et al. (2022) with primers from (Astrin & Stüben, 2008), for OPS we followed (Moškrič & Verovnik, 2019) with primers from (Audzijonyte et al., 2008), and for 28S we followed (Hou et al., 2007) with primers from the same study. The OPS marker was heterozygous as indicated by double peaks in chromatograms. The double peaks were coded using the IUPAC nucleotide ambiguity codes and haplotypes were phased using PHASE (Stephens et al., 2001) implemented in DnaSP 6 (Rozas et al., 2017). Only phased OPS haplotypes were used in subsequent analyses. Sequences were aligned using MUSCLE (Edgar, 2004) implemented in MEGA 6 (Tamura et al., 2013). The COI and OPS sequences were subsequently amino-acid translated to check for stop codons and reading frame shifts that would indicate pseudogene amplification. None were detected. All the newly obtained sequences were submitted to GanBank (COI accession numbers: ZZZ-YYY, OPS accession numbers: ZZZ-YYY, 28S accession numbers: ZZZ-YYY) (will be provided during the revision). To the final datasets we also added 22 COI sequences from a previous study (Copilaș-Ciocianu et al., 2022). See Supplementary Table S1 for further details.

### Genetic diversity and demographic analyses

To explore the spatial patterns of haplotype distributions we constructed haplotype networks for all markers using Haploviewer (Salzburger et al., 2011). As input we used maximum-likelihood (ML) trees generated for each species individually with MEGA 6. Haplotype distribution was plotted on maps using QGIS Desktop 3.22.8 (http://www.qgis.org).

Genetic diversity indices such as number of haplotypes (Hn), haplotype diversity (Hd), nucleotide diversity (Pi), and the average number of nucleotide differences (K) were calculated for each species and maker using the DNA polymorphism function in DnaSP 6. For comparative purposes, these indices were also calculated separately for the native and invaded regions for each species.

In order to test for signs of rapid demographic expansion throughout the invaded as well as native regions, we performed several demographic tests and calculated their significance: Tajima’s D (Tajima, 1989), Fu’s Fs (Fu, 1997), R2 (Ramos-Onsins & Rozas, 2002), and Raggedness statistic (Hri) (Harpending, 1994). Tajima’s D and R2 rely on the frequency of segregating sites, Fu’s Fs on haplotype distribution, while Hri measures the smoothness of the mismatch distribution (Ramos-Onsins & Rozas, 2002). All tests were calculated in DnaSP 6. Additionally, mismatch distribution analyses that examine the frequency of observed pairwise differences against an expected distribution assuming population expansion were performed in DnaSP 6.

### Hypothesis testing

To test for patterns in genetic diversity (all four indices) between invaded and native ranges (question 1) and introduced vs. self-dispersed species (question 2) we fitted a linear mixed effects model (LMEM) for each diversity metric (four for each COI and OPS), with Range (2 levels: native, invaded) and Arrival mode (2 levels: introduction, dispersal) terms as well as their interaction term as fixed factors. Species term was included as a random factor. In such a model a significant interaction term could be interpreted as a positive answer to our question 2 (difference in diversity change slopes), while a significant Range factor could be interpreted as an overall positive answer to our question 1 (reduction in diversity in the invaded range). We log-transformed the Hn, Pi, and K values for COI (the latter two – after the addition of 10^-6^ due to zeroes present) as using raw data for the LMEMs indicated significant deviations of residuals from normality and/or homoskedasticity (tested using the Shapiro-Wilk and the Levene’s tests, respectively). For the OPS metrics no data transformation was needed. *C. warpachowskyi* was excluded from the hypothesis testing based on OPS since this marker was not amplified in his species. The LMEMs were fitted and tested using the R packages *lme4* (Bates et al., 2015) and *lmerTest* (Kuznetsova et al., 2017). The models were visualized (Fig. S1) with the aid of the visreg package (Breheny & Burchett, 2017). Each LMEM was followed by four pairwise comparisons among groups using the multivariate t (mvt) P-value adjustment, implemented in the package emmeans (Lenth, 2022).

## Results

In total we obtained 360 new COI (641 bp), 154 OPS (779 bp), and 27 28S (1137 bp) sequences. Comparative mitochondrial haplotype distribution indicates a striking difference between the invaded and native regions (Table 1, Figs. 1–2). A single dominant haplotype is present in the invaded Baltic area in all species except *P. robustoides* where two dominant haplotypes are present. The main invasive haplotypes were detected in the native range for all species except *C. warpachowskyi.* Specifically, they mainly occur in the native populations of the lower Dniester and Dnieper rivers and the Dnieper-Bug estuary (Fig. 1). For *C. warpachowskyi* no samples could be obtained from the Dnieper, likely explaining why the invasive haplotype was not detected in its native range. At the nuclear level the patterns of haplotype distribution are less pronounced, without noticeable differences between the native and invaded ranges (Fig. 2). There are, however, more pronounced differences among species, some exhibiting the same dominant haplotype(s) in both ranges (e.g. *C. curvispinum, D. haemobaphes*, and *D. villosus*) while others exhibiting considerable diversity in both ranges (*O. crassus* and *P. robustoides*) (Fig. 2).

**Table 1.** Genetic diversity and tests for demographic expansion within the invaded (Baltic) and native (NE Black Sea) ranges among the deliberately introduced (INT) and self-dispersed (DIS) species.

| Species | Arrival | Range | Hn | Hd | Hd - SD | Pi | Pi - SD | K | K - variance | Tajima's D | P | Fu's Fs | P | R2 | P | Hri | P |
| --- | --- | --- | --- | --- | --- | --- | --- | --- | --- | --- | --- | --- | --- | --- | --- | --- | --- |
| COI |  |  |  |  |  |  |  |  |  |  |  |  |  |  |  |  |  |
| *C. warpachowskyi* | INT | Invaded | 2 | 0.037 | 0.035 | 0.00006 | 0.00005 | 0.037 | 0.011 | -1.095 | 0.126 | -1.701 | **0.031** | 0.135 | 0.601 | 0.859 | 1.000 |
|  |  | Native | 5 | 0.629 | 0.125 | 0.00165 | 0.00052 | 1.086 | 0.570 | -0.992 | 0.193 | -1.316 | 0.120 | 0.128 | 0.156 | 0.062 | 0.059 |
| *O. crassus* | INT | Invaded | 2 | 0.257 | 0.110 | 0.00040 | 0.00017 | 0.257 | 0.008 | -0.133 | 0.320 | 0.341 | 0.335 | 0.129 | 0.172 | 0.302 | 0.595 |
|  |  | Native | 5 | 0.752 | 0.039 | 0.00252 | 0.00019 | 1.615 | 0.056 | 1.617 | 0.944 | 0.837 | 0.706 | 0.202 | 0.961 | 0.102 | 0.333 |
| *P. robustoides* | INT | Invaded | 5 | 0.383 | 0.055 | 0.00106 | 0.00016 | 0.681 | 0.006 | -0.260 | 0.447 | -0.656 | 0.369 | 0.088 | 0.432 | 0.383 | 0.690 |
|  |  | Native | 12 | 0.894 | 0.025 | 0.02500 | 0.00114 | 6.533 | 0.557 | -0.540 | 0.341 | 0.710 | 0.649 | 0.114 | 0.541 | 0.073 | 0.784 |
| *C. curvispinum* | DIS | Invaded | 1 | 0 | 0 | 0 | 0 | 0 | 0 | N/A | N/A | N/A | N/A | N/A | N/A | N/A | N/A |
|  |  | Native | 11 | 0.888 | 0.045 | 0.01717 | 0.00277 | 11.004 | 2.253 | -0.171 | 0.483 | 1.946 | 0.805 | 0.123 | 0.509 | 0.079 | 0.916 |
| *D. haemobaphes* | DIS | Invaded | 1 | 0 | 0 | 0 | 0 | 0 | 0 | N/A | N/A | N/A | N/A | N/A | N/A | N/A | N/A |
|  |  | Native | 5 | 0.692 | 0.119 | 0.00144 | 0.00036 | 0.923 | 0.067 | -0.964 | 0.191 | -1.963 | **0.024** | 0.120 | **0.016** | 0.151 | 0.257 |
| *D. villosus* | DIS | Invaded | 1 | 0 | 0 | 0 | 0 | 0 | 0 | N/A | N/A | N/A | N/A | N/A | N/A | N/A | N/A |
|  |  | Native | 8 | 0.454 | 0.111 | 0.00109 | 0.00036 | 0.701 | 0.019 | -2.125 | **0.002** | -5.560 | **<0.0001** | 0.063 | **0.046** | 0.112 | 0.146 |
| OPS |  |  |  |  |  |  |  |  |  |  |  |  |  |  |  |  |  |
| *O. crassus* | INT | Invaded | 7 | 0.815 | 0.048 | 0.00276 | 0.00028 | 2.153 | 0.771 | 1.152 | 0.882 | -0.336 | 0.458 | 0.179 | 0.959 | 0.031 | **0.022** |
|  |  | Native | 7 | 0.354 | 0.102 | 0.0018 | 0.0008 | 1.400 | 0.493 | -2.254 | **0.002** | -1.301 | 0.235 | 0.104 | 0.346 | 0.327 | 0.981 |
| *P. robustoides* | INT | Invaded | 8 | 0.278 | 0.076 | 0.00139 | 0.00046 | 1.085 | 0.374 | -1.974 | **0.006** | -2.398 | 0.106 | 0.066 | 0.123 | 0.543 | 0.963 |
|  |  | Native | 8 | 0.627 | 0.101 | 0.00315 | 0.00068 | 2.458 | 0.880 | -0.672 | 0.270 | -0.729 | 0.377 | 0.117 | 0.411 | 0.294 | 0.999 |
| *C. curvispinum* | DIS | Invaded | 2 | 0.063 | 0.058 | 0.00008 | 0.00007 | 0.063 | 0.022 | -1.142 | 0.132 | -1.265 | 0.058 | 0.174 | 0.673 | 0.770 | 0.782 |
|  |  | Native | 2 | 0.091 | 0.081 | 0.00012 | 0.00010 | 0.091 | 0.033 | -1.162 | 0.147 | -0.957 | 0.071 | 0.208 | 0.993 | 0.678 | 1.000 |
| *D. haemobaphes* | DIS | Invaded | 3 | 0.601 | 0.080 | 0.00121 | 0.00032 | 0.941 | 0.351 | 0.219 | 0.640 | 0.980 | 0.687 | 0.157 | 0.569 | 0.118 | 0.296 |
|  |  | Native | 4 | 0.576 | 0.097 | 0.00102 | 0.00024 | 0.793 | 0.287 | -0.031 | 0.488 | -0.356 | 0.353 | 0.132 | 0.422 | 0.076 | **0.042** |
| *D. villosus* | DIS | Invaded | 2 | 0.100 | 0.088 | 0.00026 | 0.00023 | 0.200 | 0.074 | -1.513 | **0.047** | -0.025 | 0.226 | 0.218 | 0.786 | 0.830 | 0.998 |
|  |  | Native | 4 | 0.233 | 0.086 | 0.00048 | 0.00019 | 0.377 | 0.132 | -1.015 | 0.183 | -1.639 | 0.059 | 0.075 | 0.109 | 0.472 | 0.776 |
| 28S |  |  |  |  |  |  |  |  |  |  |  |  |  |  |  |  |  |
| *C. warpachowskyi* | INT | Invaded | 1 | 0 | 0 | 0 | 0 | 0 | 0 | N/A | N/A | N/A | N/A | N/A | N/A | N/A | N/A |
|  |  | Native | 1 | 0 | 0 | 0 | 0 | 0 | 0 | N/A | N/A | N/A | N/A | N/A | N/A | N/A | N/A |

Hn—haplotype number; Hd—haplotype diversity; Pi—nucleotide diversity; K—Average number of nucleotide differences; SD—standard deviation;

**Fig. 2.**
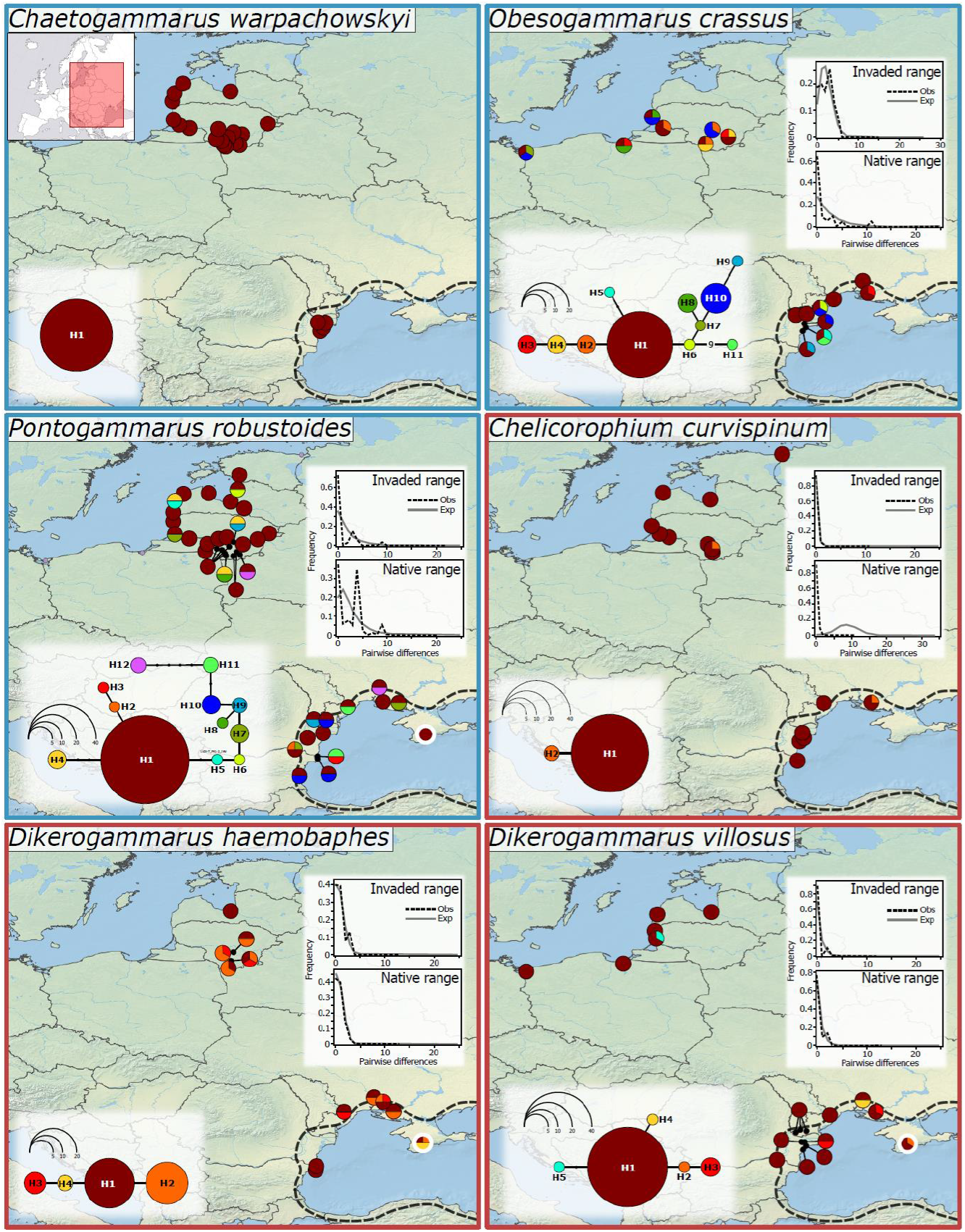
Patterns of nuclear (OPS for all species, 28S for *C. warpachowskyi*) haplotype distribution between the native and invaded ranges. Insets on the lower left depict haplotype networks while on the upper right are mismatch distributions. The native range is shown with a dashed black line. The site indicated with a thick white outline is the Simferopol WR which lies in the native range but was artificially populated. Species that were deliberately introduced and that dispersed on their own are indicated with a blue and red frame, respectively.

At the mitochondrial level, demographic tests and mismatch distribution analyses indicate demographic expansion and genetic bottleneck in the invaded Baltic region only in *C. warpachowskyi* (Fu’s Fs = −1.701, p = 0.031) (Table 1). For the species that dispersed on their own these metrics could not be calculated since only one haplotype was present (Table 1). Nevertheless, this drastic reduction of haplotype number clearly indicates a genetic bottleneck. In the native region both *Dikerogammarus* species showed signs of demographic expansion (significant Fu’s Fs and R2 tests and mismatch distribution) (Table 1). At the nuclear level, only *P. robustoides* and *D. villosus* showed signs of demographic expansion and genetic bottlenecks in the invaded Baltic region (significant Tajima’s D values). In the native range, signs of rapid expansion were observed only in *O. crassus* (Table 2).

**Table 2.** Linear mixed effects models of COI and OPS markers diversity metrics: model coefficients and analysis of variance (type III tests using Satterthwaite’s approximation for denominator degrees of freedom). Significant effects (P < 0.05) are highlighted. COI metrics Hn, Pi, and K were log-transformed. Tested terms: Arrival mode (via introduction/dispersal) and Range (native, invaded).

| Response metric | Term | COI marker |  |  | OPS marker |  |  |
| --- | --- | --- | --- | --- | --- | --- | --- |
| | | $b \pm SE$ | $F$ | $P$ | $b \pm SE$ | $F$ | $P$ |
| Hn | Arrival mode | $0.13 \pm 0.28$ | 2.8 | 0.146 | **$-4.17 \pm 0.60$** | **78.4** | **$< 0.001$** |
| | Range | **$-0.90 \pm 0.13$** | **245.0** | **$< 0.001$** | $0.00 \pm 0.45$ | 3.0 | 0.144 |
| | Arrival:Range | **$-1.13 \pm 0.19$** | **36.2** | **$< 0.001$** | $-1.00 \pm 0.58$ | 3.0 | 0.144 |
| Hd | Arrival mode | $-0.08 \pm 0.10$ | 3.0 | 0.135 | $-0.19 \pm 0.20$ | 2.2 | 0.197 |
| | Range | **$-0.53 \pm 0.07$** | **132.1** | **$< 0.001$** | $0.06 \pm 0.18$ | 0.0 | 0.966 |
| | Arrival:Range | $-0.15 \pm 0.11$ | 1.9 | 0.217 | $-0.10 \pm 0.24$ | 0.2 | 0.688 |
| Pi | Arrival mode | **$-0.45 \pm 0.85$** | **16.7** | **0.006** | **$-0.00 \pm 0.00$** | **24.4** | **$< 0.001$** |
| | Range | **$-2.77 \pm 0.57$** | **176.9** | **$< 0.001$** | $-0.00 \pm 0.00$ | 0.4 | 0.563 |
| | Arrival:Range | **$-5.24 \pm 0.81$** | **41.9** | **$< 0.001$** | $0.00 \pm 0.00$ | 0.3 | 0.606 |
| K | Arrival mode | **$-0.16 \pm 0.77$** | **86.8** | **$< 0.001$** | **$-1.51 \pm 0.39$** | **24.4** | **$< 0.001$** |
| | Range | **$-2.49 \pm 0.57$** | **441.3** | **$< 0.001$** | $-0.31 \pm 0.43$ | 0.4 | 0.564 |
| | Arrival:Range | **$-11.98 \pm 0.81$** | **220.0** | **$< 0.001$** | $0.29 \pm 0.55$ | 0.3 | 0.609 |

Hn-haplotype number; Hd-haplotype diversity; Pi-nucleotide diversity; K-Average number of nucleotide differences.

With respect to ranges, all species showed a pronounced reduction of genetic diversity at the mitochondrial but not nuclear marker in the invaded range relative to the native range (Tables 1,2, Fig. 3). The LMEMs (Table 2, Fig. S1) indicated that mitochondrial genetic diversity was generally reduced in the invaded range (significant Range effect at all metrics), but nuclear diversity was not (Table 2). Moreover, the self-dispersed species also lost more mitochondrial genetic diversity than the introduced ones (significant interaction effect at all metrics except Hd). While within the native range no differences were observed, in the invaded range the introduced species generally exhibited significantly higher mitochondrial genetic diversity than the self-dispersed species (significant pairwise comparisons within the native range group for all metrics except Hd). Interestingly, the self-dispersed species had an overall lower nuclear genetic diversity than the introduced ones (significant Arrival effect at all metrics except Hd) within both native and introduced ranges.

**Fig. 3.**
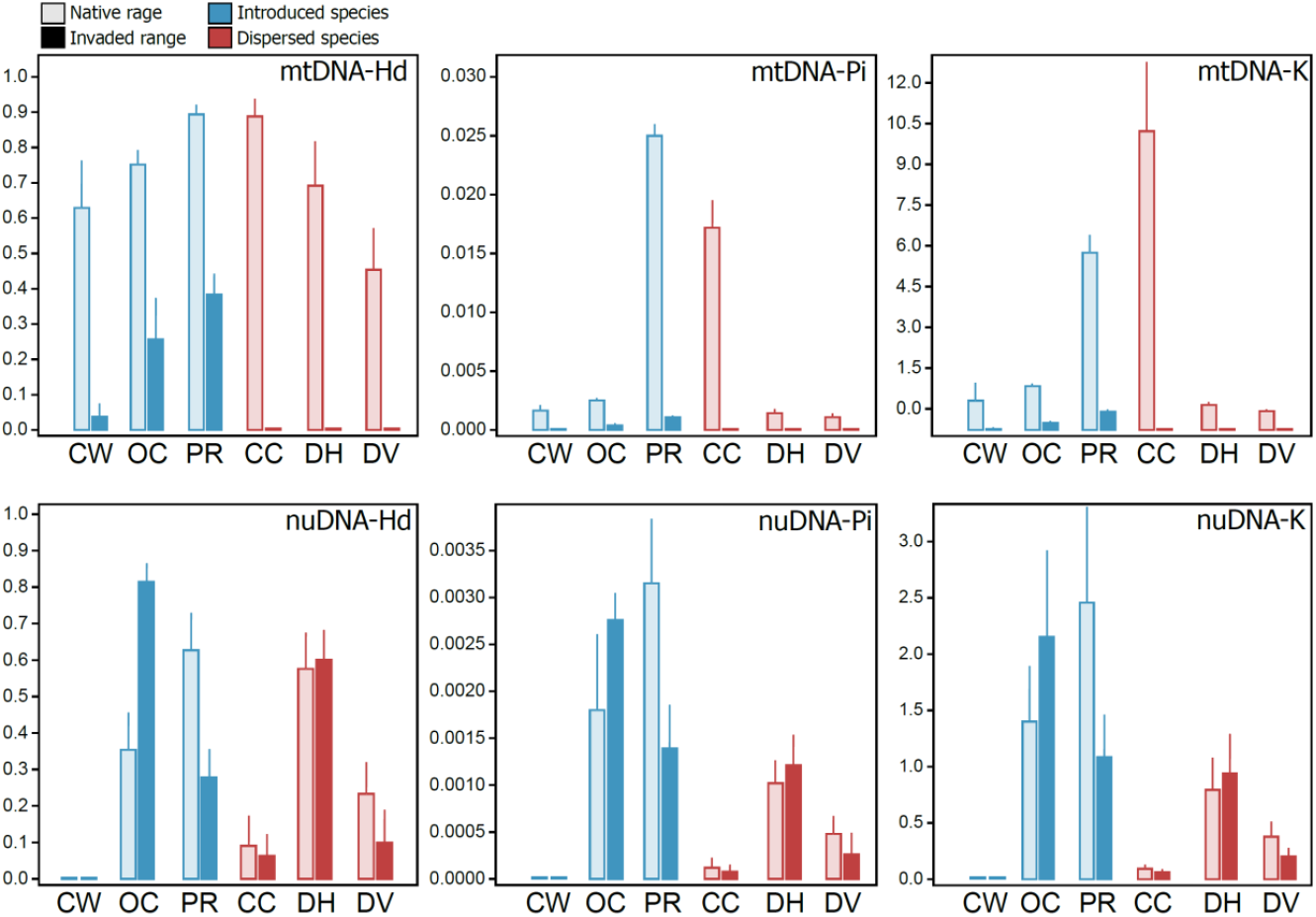
Metrics of genetic diversity for the mitochondrial (COI) marker (top) and nuclear (OPS for all species, 28S for C. warpachowskyi) marker (bottom). Metric abbreviations: Hd–haplotype diversity, Pi–nucleotide diversity, K–average number of nucleotide differences. Species abbreviations: CW–*Chaetogammarus warpachowskyi*, OC–*Obesogammarus crassus*, PR–*Pontogammarus robustoides*, CC–*Chelicorophium curvispinum*, DH–*Dikerogammarus haemobaphes*, DV–*Dikerogammarus villosus*.

## Discussion

Our results revealed surprising patterns of genetic variation in relation to geographical ranges (native vs. invaded) and arrival mode (deliberately introduced vs. self-dispersed species) to the non-native Baltic region. We find that in all six investigated species mitochondrial but not nuclear genetic diversity is reduced in the invaded range relative to the native one. Intriguingly, the deliberately introduced species exhibit higher genetic diversity in the invaded range than the species that dispersed on their own. Below we discuss these patterns in more detail and provide putative explanations.

It has long been assumed that invasive species experience a drastic reduction of genetic variation outside the native range due to bottlenecks (Baker & Stebbins, 1965). However, a plethora of studies have indicated that this is seldom the case and often alien populations have comparable genetic diversity relative to source populations due to multiple introductions and high propagule pressure (Kolbe et al., 2007; Roman & Darling, 2007; Guo et al., 2015). This pattern has been reported in many Ponto-Caspian taxa studied to date ranging from crustaceans to mollusks and fishes (Stepien et al., 2005; Audzijonyte et al., 2009, 2017; Rewicz et al., 2015; Jażdżewska et al., 2020). Our study partially confirms these findings as we detected a decrease in genetic variation at the mitochondrial but not nuclear level across all six investigated species. Such discrepancy could be due to the fact that the mitochondrial genome is haploid, uniparentally inherited and lacks recombination in amphipods and most other taxa. One the other hand, the investigated nuclear marker exhibited high levels of heterozygosity, likely reflecting the high heterozygosity and large genomes commonly encountered in amphipods (Rees et al., 2007; Kao et al., 2016; Jeffery et al., 2017). Nevertheless, given that we sequenced only one nuclear marker, these patterns should be studied further using reduced representation genomic approaches based on single nucleotide polymorphisms (SNPs) which have been proven useful in amphipods (Hupalo et al., 2022). It is likely that a genomic approach might still reveal a certain loss of nuclear genetic variation relative to the native range, but not at the same magnitude as observed for the mitochondrial genome.

Our most significant finding is that in the invaded Baltic range the deliberately introduced species have an overall higher mitochondrial and nuclear genetic diversity than the species that dispersed on their own. Interestingly, in the native range this difference persists only at the nuclear level, while mitochondrial diversity is comparable between the two groups. This discrepancy indicates that indeed introduction mode could play a role, but other factors such as species-specific genomic architecture coupled with phylogenetic effects might also be at play (see below).

It appears that introduction mode only affected mitochondrial diversity, since it differs strongly between the introduced and self-dispersed species only in the invaded range. These patterns could be explained by the fact that the introduced species were translocated in relatively large amounts (hundreds to thousands of specimens) directly by air from the Simferopol and Dnieper WRs to the Kaunas WR (Vaitonis et al., 1990), thus likely partially bypassing the initial genetic bottleneck. Moreover, after successful acclimatization in the Kaunas WR in the 1960s, tens of thousands of specimens were subsequently deliberately introduced to hundreds of waterbodies in a stepwise fashion throughout Lithuania, Latvia, Estonia, and Russia until the late 1980s (Vaitonis et al., 1990; Arbačiauskas et al., 2017). Such a pattern of introductions likely helped to quickly spread genetic diversity before being lost to genetic drift.

From an ecological point of view, the deliberately introduced species are generally more associated with lacustrine environments and have not spread as much on their own outside the native range (Copilaș-Ciocianu & Sidorov, 2022; Copilaș-Ciocianu et al., 2022b). On the contrary, the species that dispersed on their own to the Baltic region are more associated with riverine habitats and have substantially dispersed outside the native range, being among the most widespread Ponto-Caspian invaders (Rewicz et al., 2014; Copilaș-Ciocianu & Sidorov, 2022; Copilaș-Ciocianu et al., 2022b). Their affinity for flowing water suggests a superior colonization ability and higher potential for spreading via river networks and interconnecting canals. However, this colonization ability might also explain their reduced genetic diversity in the Baltic area relative to the deliberately introduced species. Given that this region represents the northern range limit of all three self-dispersed species, they may be subjected to various range margin effects such as depleted genetic variation with potential consequences on adaptive potential and persistence (Bridle & Vines, 2007; Hill et al., 2011; Takahashi et al., 2016).

The observation that the introduced species exhibit a higher nuclear genetic diversity than the self-dispersed species in both native and non-native ranges suggests that this discrepancy could be explained by species-specific genomic features and evolutionary relationships. In the related Baikal Lake radiations of gammaroidean amphipods there is an 8-fold variation in genome size among species that is positively related to depth, body size and diversification rate (Jeffery et al., 2017). Similar patterns of genomic size variation could also occur in the Ponto-Caspian taxa given that their ecological and morphological diversity is reminiscent to that of the Baikalian radiations (Copilaș-Ciocianu & Sidorov, 2022). Thus, it is likely that genomic size variation might be reflected in the observed patterns of genetic diversity among the focal species. Furthermore, taking into account phylogenetic relationships, the introduced *O. crassus* and *P. robustoides* and the dispersed *D. haemobaphes* and *D. villosus* are more related to one another than to the other species in our study (Hou et al., 2014; Copilaș-Ciocianu et al., 2022a; Morhun et al., 2022). Thus, they may share similar genomic features that could drive the observed patterns. Teasing away between the effects of evolutionary history and introduction mode on patterns of genetic diversity would require a larger dataset in terms of species and genetic data.

The well-documented introduction history of the focal taxa allows us to further test the utility of mitochondrial markers in tracing the origin of Ponto-Caspian invaders. Although these markers have proven useful in all of the crustacean species studied to date (Cristescu et al., 2001, 2004; Audzijonyte et al., 2009, 2017; Rewicz et al., 2015; Jażdżewska et al., 2020), four of the species included in our study (*C. warpachowskyi, C. curvispinum, O. crassus* and *P. robustoides*) had very limited sequence data available until now, especially from the non-native range (Cristescu & Hebert, 2005; Hou et al., 2014; Copilaș-Ciocianu et al., 2022). Here we confirm that the main invasive haplotypes (including from the Simferopol WR) can be traced to the native populations of the Dnieper-Bug estuary in all species except *C. warpachowskyi* which we did not sample from this area (possibly extinct). Unfortunately, we could not obtain specimens from the Dnieper WR where these species were also acclimatized before being introduced to Lithuania. However, given the known introduction history, these haplotypes should also be similar to the ones from the Dnieper-Bug system. Furthermore, confirming the Dnieper-Bug origin of the species that dispersed on their own to the Baltic region further emphasizes the importance of the Central Corridor (i.e. Dnieper–Vistula–Oder–Rhine and interconnecting canals) as a dispersal pathway for Ponto-Caspian fauna (Bij de Vaate et al., 2002; Copilaș-Ciocianu et al., 2022b). One remaining issue is that the rare haplotypes found at single locations in the invaded range were not detected in the native rage. Given the relatively short time since the introduction it is unlikely that these are novel variants that appeared in the invaded range. Most likely they remained undetected in the native range due to insufficient sample size or are possibly extinct there.

With respect to the native range, we find that all species except *D. villosus* exhibit a significant geographical structure of mitochondrial haplotypes with a divide between the west (Danube and surroundings) and east (Dniester and Dnieper-Bug). Although this pattern was not detected for *D. villosus*, which exhibits a single dominant haplotype throughout the entire region, it was confirmed with nuclear microsatellites (Rewicz et al., 2015). Similar patterns of differentiation across the Danube and the Dniester/Dnieper drainages have been reported for various other Ponto-Caspian crustaceans (Cristescu et al., 2001, 2004; Audzijonyte et al., 2015) and are most likely a result of the region’s geological history (Krijgsman et al., 2019).

## Conclusion

We highlight a significant loss of genetic diversity of alien Ponto-Caspian amphipods in the invaded Baltic range relative to the donor NW Black Sea range, but only at the mitochondrial level. Overall, nuclear genetic diversity did not significantly decrease in the invaded range. We also find consistent evidence that deliberately introduced species have a higher mitochondrial and nuclear genetic diversity than the species that dispersed on their own to the Baltic region. Furthermore, mitochondrial markers have once more proven useful as they correctly traced the donor populations in accordance to the known invasion history. Overall, introduction mode appears to influence genetic diversity outside the native range only at the mitochondrial level. A genomic approach coupled with a broader taxonomic coverage could provide more insight and control for phylogenetic relationships.

## Acknowledgements

We thank Jūratė Lesutienė and Gintautas Vaitonis for help during the fieldwork in Ukraine in 2012. This study was supported by the Research Council of Lithuania (Contract No. S-MIP-20-26).

## Data availability

The DNA sequences generated during this study are available in GenBank (COI: ZZZ-YYY, OPS: ZZZ-YYY, 28S: ZZZ-YYY) (accession numbers will be provided during the revision). List of sampling localities and associated information is provided in Table S1.

---

**Fig. S1.**
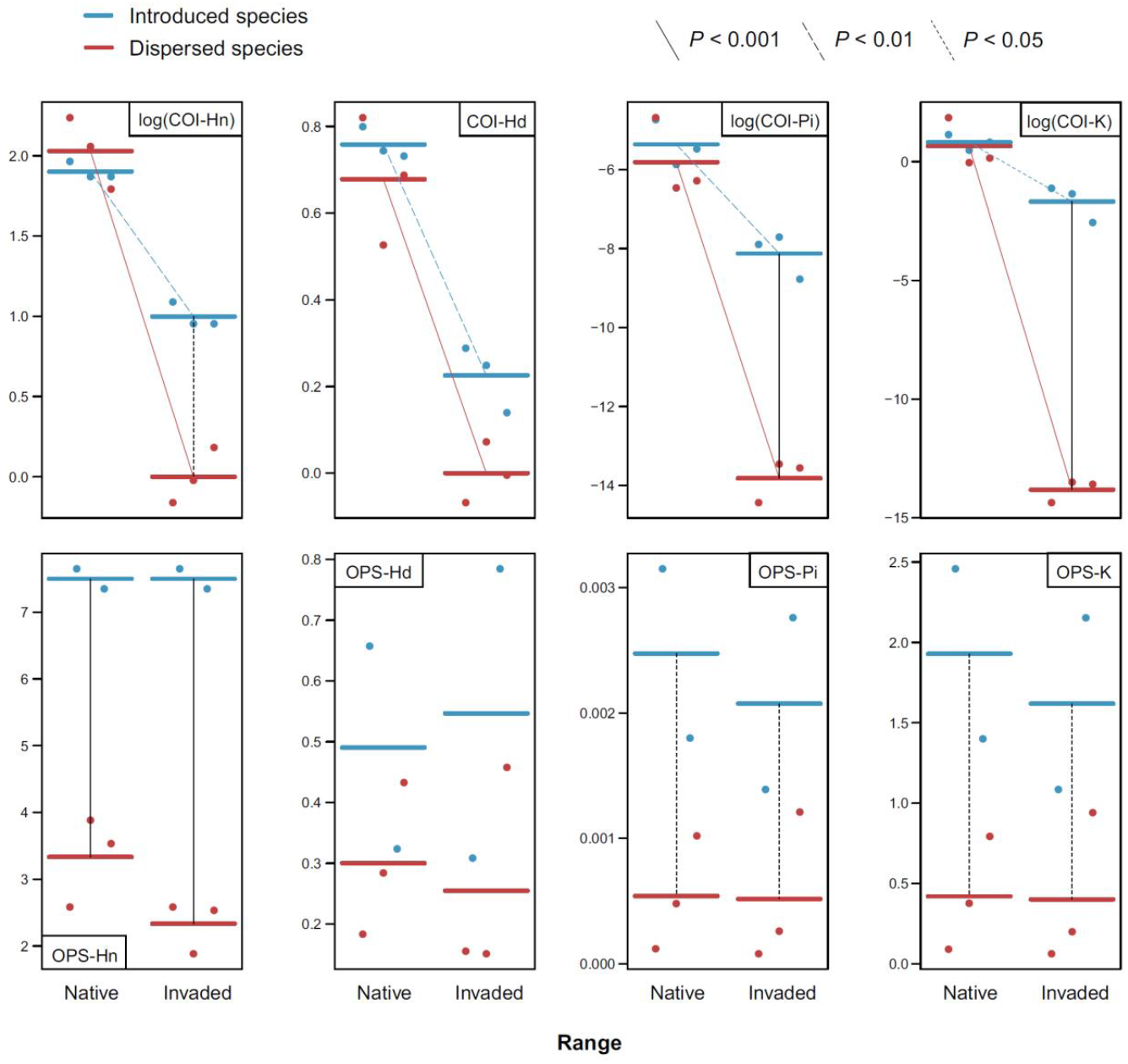
Predictions and residuals of the LMEMs for the COI (top) and OPS (bottom) marker diversity metrics (see Table 2 for model effects tests). Metric abbreviations: Hn – haplotype number, Hd – haplotype diversity, Pi – nucleotide diversity, K – average number of nucleotide differences. Thick lines represent estimated means. Significant multiple comparisons are indicated by thin connecting lines of different patterns, absence indicating results with P > 0.05.

